# The Long Noncoding RNA *Playrr* Regulates *Pitx2* Dosage and Protects Against Cardiac Arrhythmias

**DOI:** 10.1101/2022.09.20.508562

**Authors:** Frances L. Chen, Eva M. Oxford, Shao-Pei Chou, Na Li, John P. Leach, Sienna K. Perry, Bhargav D. Sanketi, Christina Cong, Sophie A. Kupiec-Weglinski, Rebecca Dubowitz, Erin Daugherity, James F. Martin, Charles G. Danko, Natasza A. Kurpios

## Abstract

**Rationale:** The most significantly associated atrial fibrillation (AF) risk loci in humans map to a noncoding gene desert upstream of the evolutionarily conserved left-right (LR) transcription factor *Pitx2*, a master regulator of LR asymmetric organ development. *Pitx2* dosage is fundamentally linked to the development of sinus node dysfunction (SND) and AF, the most common cardiac arrhythmia affecting adults, but the mechanistic basis for this remains obscure. We identified a conserved long noncoding RNA (lncRNA), *Playrr*, which is exclusively transcribed on the embryo’s right side, opposite to *Pitx2* on the left, that participates in mutually antagonistic transcriptional regulation with *Pitx2*.

**Objective:** The objective of this study was to investigate a role of *Playrr* in regulating *Pitx2* transcription and protecting against the development of cardiac rhythm disturbances.

**Methods and Results:** *Playrr* expression in the developing heart was analyzed with RNA in situ hybridization. *Playrr* was expressed asymmetrically (on the right) to *Pitx2* (on the left) in developing mouse embryos, including in mouse embryonic sinoatrial node cells. We utilized CRISPR/Cas9 genome editing in mice to target *Playrr*, generating mice lacking *Playrr* RNA transcript (*Playrr*^*Ex1sj*^ allele). Using qRT-PCR we detected upregulation of the cardiac isoform, *Pitx2c*, during visceral organ morphogenesis in *Playrr*^*Ex1sj*^ mutant embryos. Surface ECG (AliveCor®) and 24-hour telemetry ECG detected bradycardia and irregular interbeat (R-R) intervals suggestive of SND in *Playrr*^*Ex1sj*^ mutant adults. Programmed stimulation of *Playrr*^*Ex1sj*^ mutant adults resulted in pacing-induced AF. Within the right atrium of *Playrr*^*Ex1sj*^ mutant hearts, Masson’s trichrome stain revealed increased collagen deposition indicative of fibrosis, and immunofluorescence demonstrated mis-localization of Connexin 43 in atrial cardiomyocytes. These findings suggested an altered atrial substrate in *Playrr*^*Ex1sj*^ adult mice. Finally, transcriptomic analysis by chromatin run-on and sequencing (ChRO-seq) in atria of *Playrr*^*Ex1sj*^ mutant mice compared to wild type controls revealed differential expression of genes involved in cell-cell adhesion and motility, fibrosis, and dysregulation of the key cardiac genes *Tbx5* and *Hcn1*.

**Conclusions:** Adult mice lacking functional *Playrr* lncRNA transcript have baseline bradyarrhythmia and increased susceptibility to AF. These cardiac phenotypes are similar to those observed in *Pitx2* heterozygous mice. Interactions between *Pitx2* and *Playrr* may provide a genetic mechanism for modulating *Pitx2* dosage and susceptibility to SND and AF.

## Introduction

Atrial fibrillation (AF) is the most prevalent sustained cardiac arrhythmia in the human population leading to increased morbidity and mortality.^1^ Sinus node dysfunction (SND) is defined by abnormal impulse formation and propagation from the sinoatrial node (SAN) or pacemaker region, and is considered a major risk factor for the development of AF.^2^ Indeed, the two arrhythmias are interconnected: over 50% of patients with SND subsequently develop AF, while patients with AF develop subsequent pathological remodeling of the atrial substrate that is associated with SND.^3,4^ While inextricably related, the molecular and pathophysiological mechanisms linking these two arrhythmias are complex and remain unclear.

The homeodomain transcription factor *Pitx2* has been dubbed the “atrial fibrillation predisposition gene”^5^ because of its genetic linkage with AF variants clustering at the same genomic locus on chromosome 4q25.^6^ Importantly, *Pitx2* deficiency during development as well as postnatally has been linked with AF susceptibility in genetic mouse models.^7–9^ Perturbed *Pitx2* expression (both under and over-expression) has also been uncovered in atrial tissue of human patients with AF.^10,11^ *Pitx2* is a central driver of left-right (LR) asymmetric cardiac morphogenesis^12,13^ and is critical for asymmetric formation of the SAN exclusively in the right atrium. For example, in wild type (WT) mice, the left sided asymmetric *Pitx2* expression in the embryonic heart directly represses the SAN genetic program on the left, and loss of *Pitx2* in the mouse embryo results in right atrial isomerism with bilateral SAN formation.^14,15^ Interestingly, while SND is associated with myocardial remodeling in the right atrium, AF is most often associated with triggers and abnormal substrates originating from the posterior wall of left atrium and pulmonary vein,^16^ begging the question of whether transcriptional interactions modulating LR *Pitx2* expression could provide a molecular mechanism underlying the pathophysiology of these two intrinsically linked arrhythmias. Importantly, multiple GWAS have mapped the most significantly associated risk variants for AF to human 4q25 locus within the noncoding gene desert just 170 kb upstream of *Pitx2*.^*6,17–19*^ Recently, CRISPR mediated deletion of cis-regulatory elements upstream of *Pitx2* in this noncoding region led to decreased cardiac *Pitx2* expression and predisposition to AF.^20^ However, to date there are no functional studies demonstrating that long noncoding RNAs (lncRNAs) located within the *Pitx2* gene desert directly contribute to the development of *Pitx2-* associated arrhythmias, such as SND and AF.

Here we employ a CRISPR/Cas9-mediated splice mutant allele in mice (*Playrr*^*Ex1sj*^) to functionally investigate *Playrr*, a lncRNA transcript we have previously identified arising from a DNA cis-regulatory element upstream of the *Pitx2* locus.^21^ We detect upregulation of the cardiac isoform *Pitx2c* expression in *Playrr*^*Ex1sj*^ mutant embryos during visceral organ morphogenesis. We show that *Playrr* has significant expression in the developing heart and localizes its domain of expression to the right sided, embryonic SAN, opposite and asymmetric to the left sided *Pitx2*. Hypothesizing that *Playrr* plays a functional role in the modulation of *Pitx2* dosage and the susceptibility to cardiac arrhythmias, we demonstrate that *Playrr*^*Ex1sj*^ mutant mice have evidence of SND and are susceptible to pacing-induced AF, phenocopying loss-of-function *Pitx2* mouse models.^8,14^ Histological analyses reveal that *Playrr*^*Ex1sj*^ mutant mice have perturbed distribution and lateralization of Connexin 43 (Cx43), the major cardiac gap junction protein, and increased atrial fibrosis. Finally, we show via transcriptomic analyses of adult *Playrr*^*Ex1sj*^ mutant hearts that loss of *Playrr* leads to upregulation of the T-box family gene (*Tbx5)*, a key regulator of heart development,^22–24^ and *Hcn1* (Hyperpolarization Activated Cyclic Nucleotide Gated Potassium Channel 1), first identified in the SAN as a key regulator of heart rate,^25,26^ as well as calcium handling genes *Ryr2* (Ryanodine Receptor 2) and *Jph2* (Junctophilin 2) required for the generation of normal cardiac rhythm. Together our studies implicate a role for the lncRNA transcript *Playrr* located at the *Pitx2-*linked AF genomic locus in the maintenance of normal sinus rhythm and protection from atrial arrhythmias. Understanding the genetic basis of SND and AF provides a powerful framework for discovering and therapeutically targeting novel molecular mechanisms underlying cardiac arrhythmias.

## Methods

### Data Availability

Detailed descriptions of the methods are available in the Online Data Supplement. All data sets featured within this article will be made available by the corresponding author upon request.

### Animals

All experiments involving animals adhered to guidelines of the Institutional Animal Care and Use Committee of Cornell University, under the Animal Welfare Assurance on file with the Office Laboratory Animal Welfare. Mouse embryos were collected after setting up timed mating with the morning of the plug defined as E0.5 and staged at the time of isolation according to the EMAP eMouse Atlas Project, which extends original Theiler and Kaufman staging criteria.^27,28,29^ The following mouse strains and alleles were used for analyses: for WT and backcrossing: FVB/NJ (JAX® Laboratories (Stock No: 001800); for *Pitx2* mutants: *Playrr*^*Ex1sj 21*^; *Hcn4-CatCH-IRES-LacZ* ^*30*^; Cornell Heart Lung and Blood Resource for Optogenetic Signaling, CHROMus; JAX® Laboratories (Stock No: 033344).

### RNA in situ hybridization

E10.5 whole mouse embryos were isolated, fixed in 4% PFA/PBS O/N, dehydrated, and stored in 100% methanol prior to processing. Whole mount ISH was performed on E10.5 whole embryos, micro-dissected embryonic hearts, and 250 μm tissue chopped slices following standard protocols as previously described.^31^

### RNAScope in situ hybridization

E10.5 whole mouse embryos were isolated, fixed in 4% PFA/PBS overnight, infiltrated with 30% sucrose, and processed for OCT embedding and sectioning as previously described.^31^ 10 μm cryosections were prepared and stored in -80 °C overnight. Multiplex fluorescent RNAScope-ISH was performed according to standard ACD protocols. The following probes were commercially available or were designed as custom probes through ACD: *Playrr:* custom 20ZZ probe named Mm-D030025E07Rik targeting nucleotides 2-946 of NR_045704.1; *Shox2:* 20ZZ probe targeting nucleotides 539-1478 of NM_013665.1.

### Quantitative reverse-transcriptase PCR (qRT-PCR)

Embryonic tissue was isolated in cold PBS and stored in RNAlater. RNA was extracted with a Qiagen RNeasy miniprep kit. 2 mg of RNA was reverse transcribed using the ABI high-capacity cDNA archive kit and diluted to 20 ng/μl. For micro-dissected tissue (heart, limb buds, gut tube) from individual embryos at E10.5, TaqMan Cells to Ct Kit was adapted for low input tissues as previously described.^32–34^

The following TaqMan gene expression assays were used for relative quantification using an ABI7500 real time PCR system: Actb, Mm00607939_s1; D030025E07Rik (Playrr), Mm03937997_m1; GAPDH, Mm99999915_g1; Pitx2ab, Mm00660192_g1; Pitx2c, Mm00440826_m1; pan-Pitx2, Mm01316994_m1. Statistical analysis was performed according to MIQE guidelines, using Welch t-tests (GraphPad PRISM 9).

### AliveCor® (Kardia) Surface ECG

Three to 7-month-old mice were humanely restrained according to standard IACUC approved procedures and prepped for 30-seconds of non-invasive surface ECG screening with AliveCor® (Kardia). A minimal amount of 70% EtOH was applied to a Kimwipe™ and used to clean and expose the skin of the ventrum of each mouse. Approximately 0.5 mL of ultrasound gel was applied to the thoracic region at the level of the sternum and each mouse to establish contact with the device. AliveCor® ECGs were analyzed manually for calculation of HR and interbeat (R-R) intervals with genotypes blinded.

### Telemetry ECG on unstressed mice

7-month-old *Playrr*^Ex1sj^ mutant mice and WT with littermate controls were implanted with telemetry implant (HD-X11, Data Sciences International) following the protocol of the manufacturer. Transmitters were placed subcutaneously and ECG leads were placed in a lead II configuration with subcutaneous fixation of leads within the right and left axial areas. After a postoperative recovery period of ten days, telemetric ECG data were collected by DSI Telemetric Physiological Monitor System RPC-1 with a sampling bandwidth range of 0.1 to 200 Hz and processed by Dataquest A.R.T 4.31. ECG data was collected via Dataquest A.R.T 4.31 Acquisition software recorded data for 24 to 50-hour periods, including 12-hour light (low activity) and 12-hour dark (high activity) cycles. Mice were not stressed from handling or anesthesia during the entire length of recording, which allowed for true resting and active recordings.

### Telemetry statistical analysis

Interbeat (R-R) interval series and heart rate series statistics for mean and standard deviation of average heart rate and average R-R intervals were generated using Dataquest A.R.T 4.31 Analysis software. Mean and standard deviation of average heart rates and average R-R intervals over 10 second intervals in a 12-hour light or 12-hour dark period were compared for statistical significance using two tailed t-tests in JMP^®^ statistical software (SAS). Tachograms (R-R time series over 10 second intervals) for each 12-hour light (low activity) or 12-hour dark (high activity) cycle were visualized in R and plotted in GraphPad PRISM 7.

### ECG and Heart Rate Variability Analysis

#### Inspection of Circadian Rhythm

24-hour ECG recordings were inspected in Dataquest A.R.T 4.31 for presence of normal circadian rhythm; HR, temperature, and activity were confirmed to be higher during dark photoperiod/high-activity phase as compared to light photoperiod/low-activity phase.

#### Beat Classification and ECG analysis

Standard ECG analysis was performed with LabChart Pro v8.0 software, using the Hamilton-Tompkins QRS detection algorithm^35^ for detection of R peaks. Beat calling was performed with the following LabChart Pro v8.0 software settings for mouse-specific beat detection consistent with previous reports on ECG parameters in adult FVB mice^37,38^ : QRS width = 10 ms, R-R interval = 20 ms, pre-P baseline = 10 ms, maximum PR = 50 ms, maximum RT interval = 40 ms, ST height measured at 10 ms from alignment. The QT interval was corrected using the Bazzett formula (QTc) and the end of the T wave was estimated using the intersection of the ECG signal with the isoelectric line (rodent T wave). 12-hour light and dark periods of all mice were manually evaluated by a board-certified veterinary cardiologist (EMO) to confirm the accuracy of LabChart Pro beat-to-beat labelling. Additionally, the ECG recordings were evaluated for arrhythmias including ventricular ectopy, atrioventricular block, sinus pauses/arrest, and supraventricular ectopy. Analysis was performed by the veterinary cardiologist with genotypes of mice blinded. The LabChart Pro software ECG analysis module beat classifier view was used to pinpoint areas to manually screen for distinguishing between artifact and arrhythmia, and to further identify ectopic beats and sinus pauses for manual removal. Beat by beat R-R intervals and corresponding HR, PR interval, QRS interval, P duration and P amplitude ECG parameters were exported from LabChart Pro. Filtering of abnormal beats was performed in accordance with standard protocols using a 95% confidence interval (mean +/-2 s.d.) in R.^36^ Data visualization (frequency histograms) and statistical analysis comparing *Playrr*^Ex1sj^ WT and mutant mice of all ECG parameters was performed for all ECG parameters in the R statistical environment and JMP Student Edition v12.1 statistical software. ECG parameter means for *Playrr*^Ex1sj^ WT and mutant mice were compared using two-tailed t-tests. Statistically significant results were plotted for display in GraphPad PRISM 7.0. Condensed R-R time series (tachogram) for each photoperiod was generated by plotting filtered mean R-R intervals over every 10 seconds in GraphPad PRISM 7.0.

#### Poincaré plots

Poincaré plots were utilized in order to further characterize periods of sinus dysrhythmia in *Playrr*^Ex1sj^ mutant mice compared with WT controls. Plots were created as previously described^37,38^ by selecting ∼5,000 consecutive data points within an episode of sinus dysrhythmia and compared with segments of similar heart rate. Data points were plotted as RR intervals along the x axis and RR+1 along the y axis.

### Programmed Intra-cardiac Pacing

Programmed intracardiac pacing was performed as previously described^39^ to determine AF susceptibility. Briefly, the recordings of surface ECG and intracardiac electrograms were performed in anesthetized adult mice via right jugular vein catheterization. The mice were subjected to an overdrive pacing protocol three times. AF is defined as the occurrence of rapid, fragmented atrial electrograms with irregular R-R intervals lasting at least 1 second in length following the burst protocol. A mouse was considered as AF positive with at least two of the three pacing protocols demonstrating AF. Incidence of inducible AF for each mouse line was calculated as the percentage of AF-positive mice divided by total mice studied. AF burden is calculated by averaging the length of AF following each of the three pacing trials. During these experiments and subsequent interpretation of AF inducibility, the operator was blinded to genotype. C57Bl/6 mice were used as control mice. To test statistical significance, One-Sided Fisher’s exact test was performed for Pacing AF Incidence, and Kruskal-Wallis test with Dunn’s multiple comparison post-test for AF Burden, respectively.

### Cx43 immunofluorescence

#### Sample preparation

Left and right atrial samples were obtained from 5 WT and 9 *Playrr*^Ex1sj^ mutant mice. Median age of WT mice was 36 weeks (range 10-36 weeks), and of mutant mice was 23 weeks (range 10-36 weeks). Tissues from all mice were harvested after experiments were performed. Samples were fixed for 48-72 hours in 10% formalin and processed for paraffin embedding. Sections were processed for immunofluorescence as previously described ^40^). Briefly, sections were deparaffinized and rehydrated by stepwise immersion in xylene, 100% ethanol, 75% ethanol, 50% ethanol, and finally phosphate buffered saline (PBS) with 0.5% tween (PBS-T). Sections were rinsed for 5 minutes in ddH2O and blocked with a solution containing 5% bovine serum albumin (BSA) in PBS-T for 2 hours at room temperature. Samples were incubated with a mouse monoclonal antibody recognizing the C-terminal end of connexin 43 (Cx43) (Sigma Aldrich, Cat# MABT901) at a 1:100 dilution in 5% BSA-PBS-T overnight at 4°C. The following day, samples were rinsed with PBS and incubated with Alexa Fluor 594 anti-mouse secondary antibody (ThermoFisher Scientific) and Hoescht dye (ThermoFisher Scientific) as a nuclear marker; each diluted 1:500 in 5% BSA-PBS-T for 2 hours at room temperature. Samples were rinsed with PBS and mounted using Permafluor aqueous mounting medium (Thermo Fisher Scientific).

#### Cx43 localization microscopy and analysis

Confocal microscopy was performed using a Zeiss AxioObserver LSM 510 confocal microscope equipped with a 63X water immersion objective. Sample genotypes were masked to the operator. For each sample of left atria and right atria, between 1-7 (average 3) separate areas were photographed to represent Cx43 localization. Areas were chosen based on good longitudinal orientation of myofibrils with strong nuclear signals.

Images were analyzed using ImageJ software (v1.52). Analysis was completed by 2 investigators; one (Joseph Choi) was blinded to the genotype of the samples. Three different localization patterns for Cx43 were recognized: perpendicular to the myofiber in the area of the intercalated disc (ID), parallel to the myofiber (lateralization) or diffuse throughout the cell (DIF). These patterns were quantified for each image and normalized to the estimated number of cells per field. Numbers of cells per image were estimated by measuring the total length of all myofibrils within the image and dividing by 100 (the average atrial myocyte length was estimated to be 100μm).^41^ The difference between group means and the ratio of DIF/ID signal compared between WT and *Playrr*^Ex1sj^ mutants was tested for statistical significance using Welch’s t-test, assuming unequal variance for each group (alpha = 0.05) (GraphPad PRISM 9).

### Masson’s trichrome staining

#### Sample preparation

Masson’s trichrome stain was used to evaluate collagen deposition in 4 WT and 4 *Playrr*^Ex1sj^ mutants. One WT mouse was 40 weeks old, the remaining three were 36 weeksold (average age 37 weeks). All mutants were 36 weeks old. Left and right atrial samples were obtained and prepared as described above. Four μm thick sections were deparaffinized and were treated with Masson’s trichrome stain via standard practices utilized by the Cornell University Animal Health Diagnostic Center histopathology facility.

#### Masson’s trichrome analysis

The genotypes of the mice were masked from the researchers during analysis. Using 40X magnification, three representative images were obtained from both the right and left atria of each sample. Images were evaluated using the following protocol in ImageJ (v1.52)^42^: Under the *Image* tab, *RGB stack* was selected, then *Make Montage* was selected. Next, under the *Image* tab, *Adjust Threshold* was selected, and the threshold was set so that all areas stained blue (for collagen) were highlighted. Under the *Analyze* tab, *Set Measurements* window was opened. Within the *Set Measurements* window, *Area, Percent Area, Limit to Threshold, and Display label* were checked. Finally, under the *Analyze* tab, *Measure* was selected, and the values recorded. Percent collagen area was compared between WT RA and *Playrr*^Ex1sj^ mutants RA, and WT LA and *Playrr*^Ex1sj^ mutants LA groups. The difference between group means was tested for statistical significance using Welch’s t-test, which does not assume equal variance for each group (alpha = 0.05) (GraphPad PRISM 9).

### LacZ tissue staining

*Playrr*^Ex1sj^ mutants (*Playrr*^*Ex1sj*^*/Playrr*^*Ex1sj*^) were crossed with hemizygous *Hcn4-CatCH-IRES-LacZ/+ mice to generate Playrr*^*Ex1sj*^*/+; Hcn4-CatCH-IRES-LacZ/+* mice. This F1 generation was further crossed to generate F2 *Playrr*^*Ex1sj*^*/+;Hcn4-CatCH-IRES-LacZ/+* and *+/+;Hcn4-CatCH-IRES-LacZ/+* embryos. Embryos were isolated at E10.5, fixed in fresh 4% PFA/PBS for 1 hour at 4 °C, rinsed in rinse buffer (100 mM sodium phosphate (pH 7.3), 2 mM MgCl2, 0.01% sodium deoxycholate, 0.02% NP-40) three times for 30 min at room temperature. Tissues were stained with staining solution (rinse buffer, 5 mM potassium ferricyanide, 5 mM potassium ferrocyanide, 1 mg/mL X-gal) at room temperature for 16 hours. Samples were post-fixed overnight in 4% PFA at 4 degrees C, then transferred to sucrose/OCT for OCT embedding and sectioning as previously described.^31^

### Chromatin Run-On Sequencing (ChRO-seq)

ChRO-seq was performed as previously described^43^ on pooled left and right atrial tissues from adult 24–32-week-old WT (n=4) and *Playrr*^*Ex1sj*^ mutant mice (n=4). Reads were mapped with BWA^44^ to the *Mus musculus* reference genome (*mm10*). Mapped reads were converted to bigWig format using BedTools^45^ and the bedGraphToBigWig program in the Kent Source software package.^46^ Downstream data analysis was performed using the bigWig software package. 52,249 GENCODE (vM20) - annotated transcripts best supported from ChRO-seq data were selected using tuSelector.^47^ Analyses were limited to 42,552 gene annotations longer than 500 bp in length. Differentially expressed genes were identified using DESeq2^48^ at a false discovery rate (FDR) less than 1%. dREG was used as previously described to identify transcriptional regulatory elements.^49^ tfTarget was used to identify transcription factor motifs in TREs showing differential expression between *Playrr*^*Ex1sj*^ mutant and WT. Gene Ontology analysis was performed. Genome wide transcriptional profiles of *Playrr*^*Ex1sj*^ mutant and WT were compared using hierarchical cluster and principal component analysis.

## Results

### Loss of *Playrr* transcript results in isoform-specific upregulation of *Pitx2c* transcription in developing mouse embryos

We previously discovered a conserved lncRNA we named *Playrr* that is transcribed from a large intergenic gene desert located approximately 968 kb upstream the *Pitx2* locus, on mouse chromosome 3 (**Fig. 1A**).^21^ During organogenesis, this genomic locus participates in long-range chromatin interactions with the *Pitx2* locus, establishing LR asymmetric chromatin architecture that mirrors LR gene expression.^21^ Our previous work demonstrates that loss of *Playrr* leads to upregulation of pan-*Pitx2* expression in developing visceral organs, suggesting mutual antagonism between *Pitx2* and *Playrr* expression.^21^ Moreover, recent work demonstrates that additional long-range chromatin interactions have a direct role in noncoding regulation of *Pitx2* in preventing susceptibility to AF.^20^ Therefore, we sought to investigate transcriptional and functional roles for *Playrr* in the context of cardiac development and disease using mice lacking the *Playrr* transcript (*Playrr*^*Ex1sj*^ mutant allele). Importantly, this allele leads to a loss of *Playrr* lncRNA transcript while leaving binding sites in the associated DNA enhancer (e926) intact,^21^ and allows parsing of the function of the lncRNA transcript and its transcription from its associated DNA regulatory element (**Fig. 1B**).

**Fig. 1.**
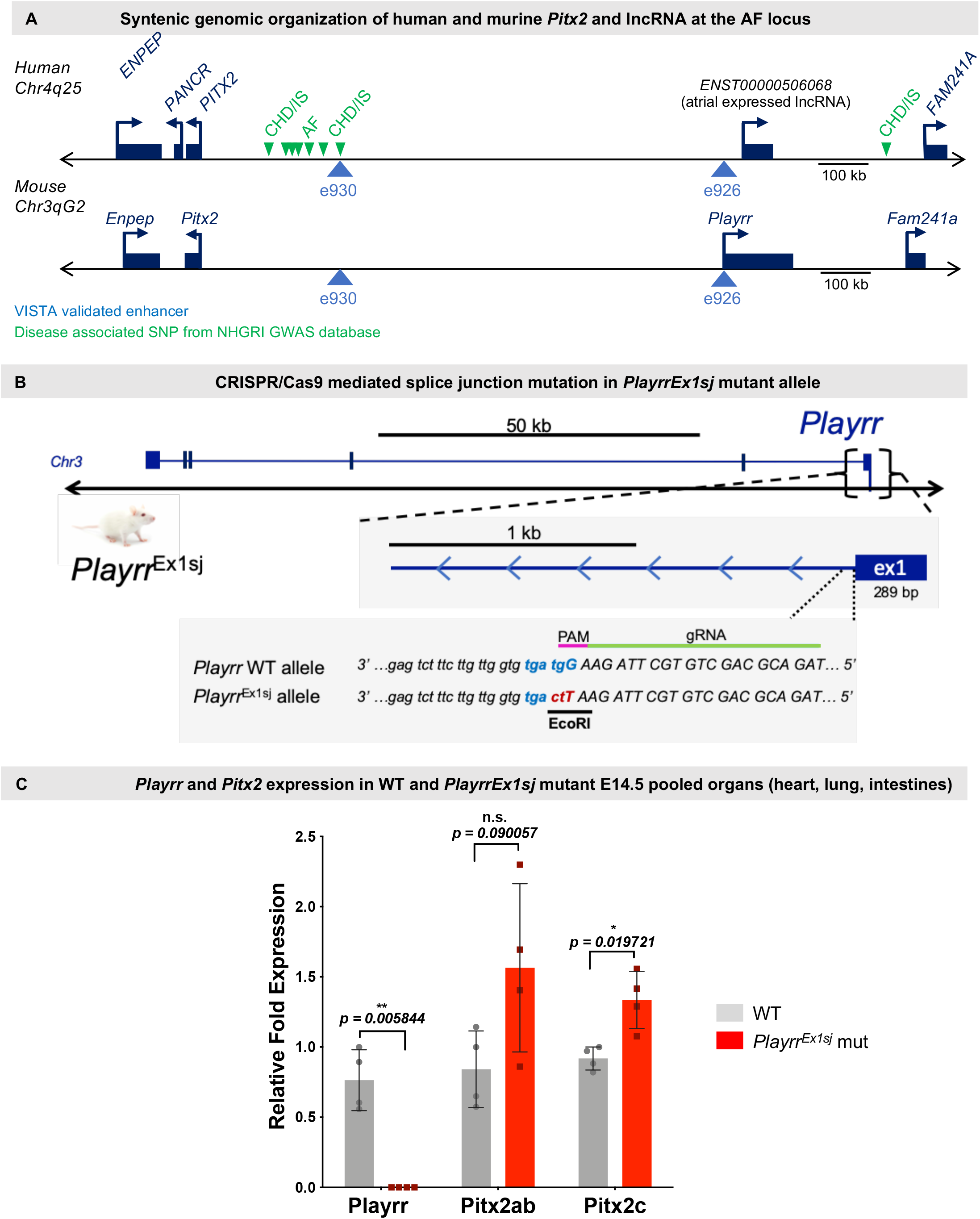
Structure of the AF risk locus, lncRNA *Playrr*^*Ex1sj*^ mutation, and *Pitx2* expression in *Playrr*^*Ex1sj*^ mice during organ development. **(A)** Structure of the atrial fibrillation locus and the genomic relationship of *Pitx2* and *Playrr*. Blue triangles denote evolutionarily conserved human/mouse VISTA enhancer sequences able to drive transgene reporter activity in mouse tissue in vivo. Green arrowheads denote GWAS implicated disease SNPs (CHD/IS = coronary heart disease/ischemic stroke; AF = atrial fibrillation). All annotations were taken from UCSC Genome Browser (Kuhn et al. 2013; Haeussler et al. 2019). **(B)** Schematic of *Playrr* splice junction mutation allele used to generate *Playrr*^*Ex1sj*^ mutant mice. **(C)** *Playrr* and *Pitx2* isoform expression assayed by qRT-PCR in E14.5 pooled organ viscera (heart, lungs, intestines) of *Playrr*^*Ex1sj*^ mutant and WT mice.

First, we confirmed loss of detectable *Playrr* RNA by qRT-PCR in *Playrr*^*Ex1sj*^ mutant mice at E10.5 and E14.5 (**Fig. S1A**, p < 0.0001). Consistent with mutually antagonistic relationship between *Pitx2* and *Playrr* expression (Welsh et al., 2015), we demonstrate upregulation specifically for the critical cardiac *Pitx2* isoform, *Pitx2c*, in *Playrr*^*Ex1sj*^ mutant mice during organ morphogenesis and embryonic (E) development (E14.5, ∼ 50% upregulation) (**Fig. 1D;** p=0.00911). Thus, loss of *Playrr* transcription results in increased expression of *Pitx2c*.

### *Playrr* is expressed in the developing SAN opposite and asymmetric to *Pitx2*

In order to investigate *Playrr* functional domains in the heart, we used qRT-PCR and RNA in situ hybridization (ISH) to assay for *Playrr* expression throughout mouse cardiac development. Using qRT-PCR, we quantified *Playrr* expression during LR patterning of the heart at E10.5, and at E14.5 when LR atrial development is complete (Krishnan et al., 2014). We found that *Playrr* expression is enriched in the heart when compared to whole embryos at E10.5 (16.9-fold) (**Fig. 2A**). Importantly, we found that *Playrr* is expressed at E10.5 in the right-sided SAN domain at the junction of the right sinus venosus and primitive right atrium (**Fig. 2B-2C)**. During development, *Pitx2c* expression inhibits SAN development on the left in a dose dependent manner and myocardial Pitx2c mutant and *Pitx2c* heterozygous embryos display intermediate properties of bilateral SAN function, highlighting that *Pitx2* dosage is critical to proper SAN patterning.^9,14^ The presence of *Playrr* within the right SAN domain during development suggests that *Playrr* also plays a role in SAN development. To validate *Playrr* expression in SAN pacemaker cells, we co-localized *Playrr* expression with the SAN marker *Shox2* using RNAScope-ISH in E10.5 WT embryos (**Fig. 2D)**. These data revealed that *Playrr* is expressed in *Shox2-*expressing SAN cells (**Fig. 2D**). Collectively, *Playrr* is highly expressed in cardiac developmental domains opposite and asymmetric to *Pitx2*, including in SAN pacemaker cells.

**Fig. 2.**
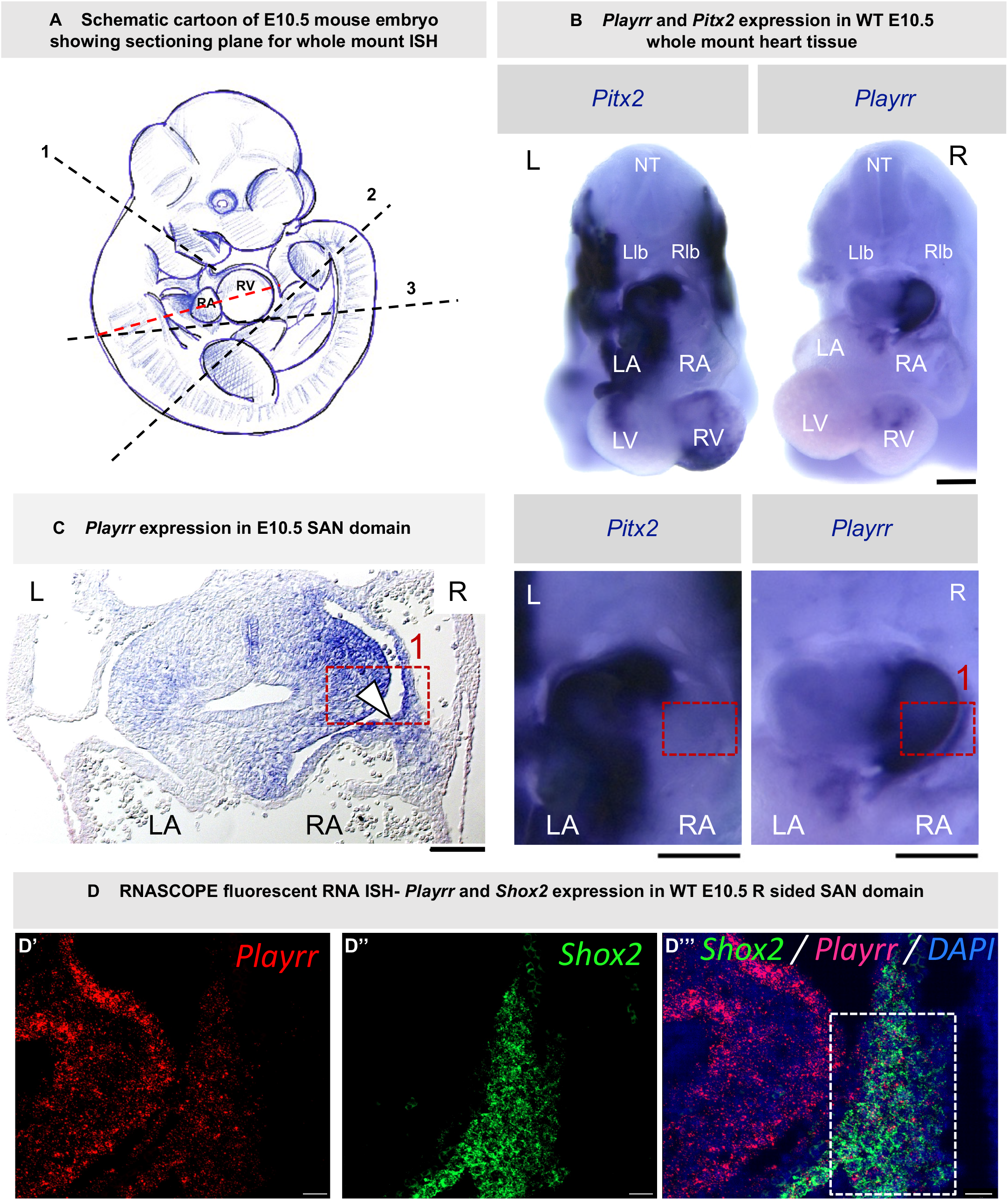
*Playrr* is expressed in the heart at E10.5 in the SAN domain complementary and asymmetric to *Pitx2*. **(A)** Cartoon of E10.5 mouse embryo showing microdissection planes for whole mount ISH shown in (B) (1, 2, 3) and plane of section ISH shown in C and D (red dotted line). **(B)** Whole mount ISH of asymmetric and complementary *Pitx2* and *Playrr* expression at E10.5 in the sinoatrial node (SAN) domain. NT (neural tube); Llb (left lung bud); Rlb (right lung bud); LA (left atrium); RA (right atrium); LV (left ventricle); RV (right ventricle). Scale bars = 2000 μm **(C)** Sectioned whole mount ISH from (B) showing *Playrr* expression in E10.5 right sided SAN domain. Dashed line box highlights area of right sinuatrial junction (1); white arrowhead = E10.5 SAN domain; LA (left atrium); RA (right atrium). Scale bar = 100 μm **(D)** RNASCOPE fluorescent ISH in right sided SAN domain at E10.5: (**D’**) *Playrr* expression (red) (**D’’**) *Shox2* expression (green) (**D’’’**) Merged DAPI (blue), *Playrr* (pink), and Shox2 (green) channels. Scale bar = 30 μm

### Adult *Playrr*^*Ex1sj*^ mutant mice display cardiac arrhythmias during restraint and at rest

Given that *Playrr* is expressed in the critical SAN spatiotemporal domain, we sought to investigate whether loss of *Playrr* affects cardiac conduction in adult mice. To efficiently and non-invasively screen *Playrr*^*Ex1sj*^ mutant mice for cardiac arrhythmias, we adapted a veterinary AliveCor^®^ ECG device for use in adult laboratory mice^50^ (**Fig. 3A)**. AliveCor^®^ surface ECG screening revealed that awake and restrained 28 week old *Playrr*^*Ex1sj*^ mutant mice had significant bradycardia compared to age-matched WT and *Playrr*^*Ex1sj*^ heterozygote controls (WT mean HR ±SD: 672±110; *Playrr*^*Ex1sj*^ heterozygote mean HR±SD=639 ±65; *Playrr*^*Ex1sj*^ mutant mean HR±SD= 384.2±132 bpm) (**Fig. 3B, C**; WT, n=2 vs *Playrr*^*Ex1sj*^ mutant, n=6; p=0.0337; *Playrr*^*Ex1sj*^ heterozygote, n=6 vs *Playrr*^*Ex1sj*^ mutant, n=6; p=0.00171). We also observed irregular R-R intervals in 5/6 *Playrr*^*Ex1sj*^ mutant mice vs 0/6 age-matched *Playrr*^*Ex1sj*^ heterozygous controls (**Fig. 3D;** *Playrr*^*Ex1sj*^ heterozygous, n=0/6; *Playrr*^*Ex1sj*^ mutant, n=5/6). Both bradycardia and irregular R-R intervals are diagnostic criteria of SND, demonstrating that *Playrr*^*Ex1sj*^ mutant mice display abnormal cardiac rhythm when restrained (**Fig. 3A-3D**).

**Fig. 3.**
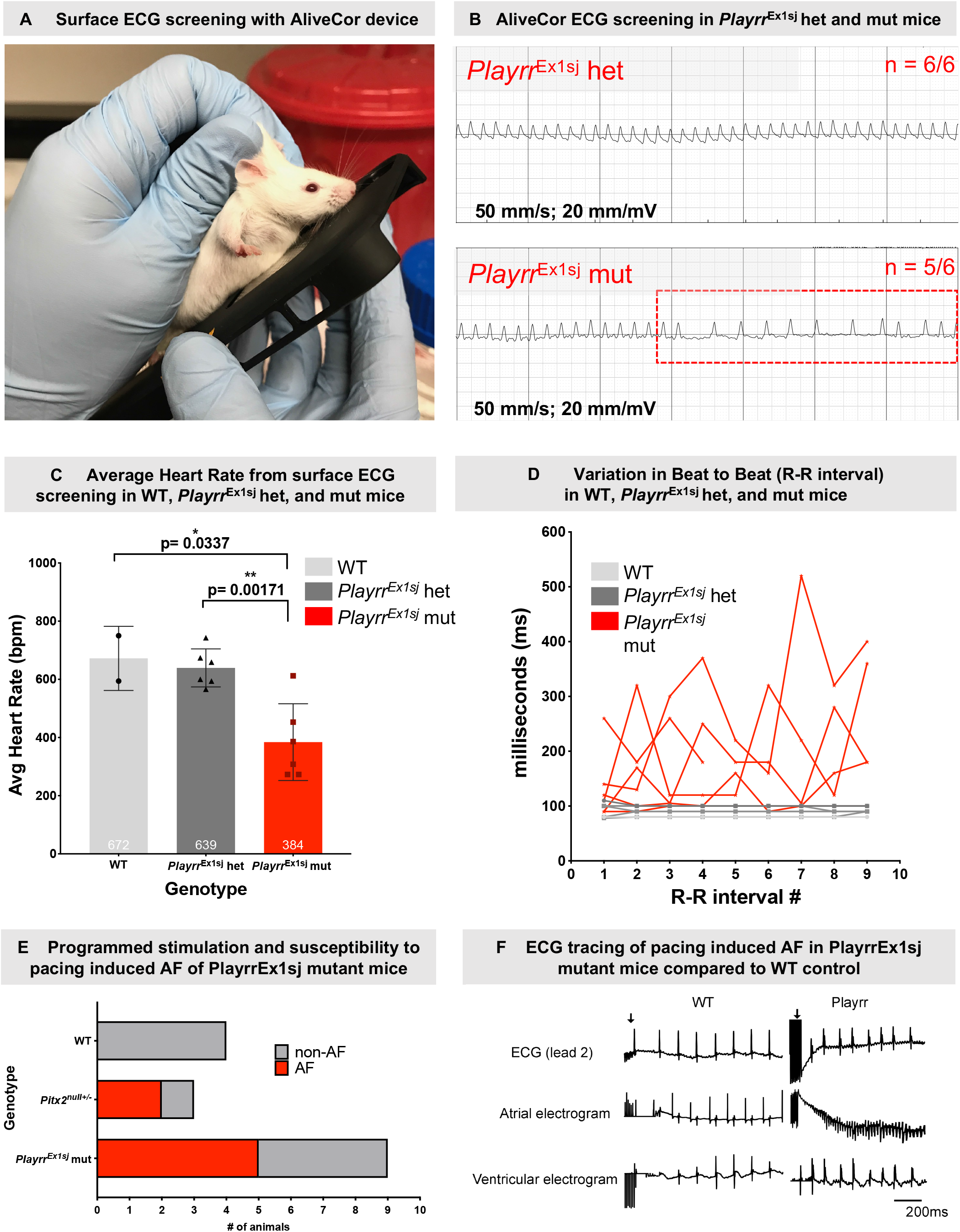
AliveCor surface ECG screening in awake, restraint-induced stressed *Playrr*^*Ex1sj*^ mice. **(A)** Photograph of surface ECG device and screening of awake, restrained mice. **(B)** ECG tracing from AliveCor screening in representative *Playrr*^*Ex1sj*^ het control (top) and mutant (bottom). **(C)** Calculated average heart rate (HR) (bpm) in *Playrr*^*Ex1sj*^ mice. WT (n=2) mean +/-SD = 672 +/-110; *Playrr*^*Ex1sj*^ het (n=6) mean +/-SD = 639 +/-65; *Playrr*^*Ex1sj*^ mutant (n=6) mean +/-SD = 384.2132 bpm. **(D)** Variation in beat to beat (R-R) interval in *Playrr*^*Ex1sj*^ mutants (grey lines) vs aged and age matched littermate WT and het controls (darker lines). **(E)** Bar charts showing incidence (red) and non-incidence (grey) of pacing induced AF following programmed stimulation in WT (n=0/4), positive control heterozygous *Pitx2*^*null/+*^ mutants (n=2/3), and *Playrr*^*Ex1sj*^ mutants (n=5/9). **(F)** Representative ECG tracings showing lead 2 surface ECG (top) and intracardiac leads atrial (middle), and ventricular (bottom) electrograms in WT (left) and *Playrr*^*Ex1sj*^ mutant.

Based on the association of *Playrr* with *Pitx2* at the AF locus, we sought to determine if *Playrr*^*Ex1sj*^ mutant mice exhibit susceptibility to pacing-induced AF. Pacing-induced AF has been previously demonstrated in *Pitx2* heterozygous (*Pitx2*^null+/-^) and in cardiac-specific compound *miR-17-92* and *miR-106b-25* haploinsufficient mice.^8,9^ Remarkably, we found that approximately 45% of *Playrr*^*Ex1sj*^ mutant mice are susceptible to pacing-induced AF compared to WT controls, where 0% of mice subjected to programmed stimulation developed AF. This is comparable to *Pitx2*^null+/-^ positive controls, where 67% demonstrate pacing-induced AF.(**Fig. 3E**; WT, n=0/4 vs *Pitx2*^null+/-^ n=2/3 vs *Playrr*^*Ex1sj*^ mutant, n=5/9).

Next, we sought to evaluate the cardiac rate and rhythm in unrestrained *Playrr*^*Ex1sj*^ mutant 28 week old adult mice over a longer timeframe. Using 24-hour telemetry ECG we found baseline arrhythmias in *Playrr*^*Ex1sj*^ mutant mice that were not observed in WT controls **(Fig. 4A’)** including increased and irregular R-R intervals and paroxysmal episodes of sinus dysrhythmia (**Fig. 4A”-A’’’**). Analysis of summary ECG parameters by 12-hour activity/photoperiod of *Playrr*^*Ex1sj*^ mutant mice revealed significant decrease in the average heart rate compared to WT controls (including littermates and non-littermates) **(Fig. 4B**; WT, n=6 vs mutant, n=8; light: p=0.0367; dark: p=0.0356). 24-hour tachograms revealed increased R-R interval variability compared to WT controls (n=6) in both 12-hour light (**Fig. 4C**; WT, n=6 vs *Playrr*^*Ex1sj*^ mutant, n=8) and 12-hour dark (**Fig. 4D)** photoperiods, with no statistically significant differences in PR interval, P duration, P amplitude, or PR interval (data not shown). Poincaré plots of 5000 consecutive beats in *Playrr*^*Ex1sj*^ mutants demonstrated loss of normal ellipsoid shape and notably altered beat-to-beat variation compared with rate matched intervals in controls (**Fig. 4E**; WT, n=6 vs *Playrr*^*Ex1sj*^ mutant, n=8). Collectively, these findings reveal that *Playrr*^*Ex1sj*^ mutant mice have characteristics of SND, defined by bradycardia and episodes of irregular R-R intervals with normal PR intervals. Interestingly, not only is SND a risk factor for AF in human patients, but the presence of SND in *Playrr*^*Ex1sj*^ mutant mice mirrors the phenotype of mice with cardiac-specific inactivation of the *miR-17-92* and *miR-106b-25* pathways downstream of Pitx2.^16,51^

**Fig. 4:**
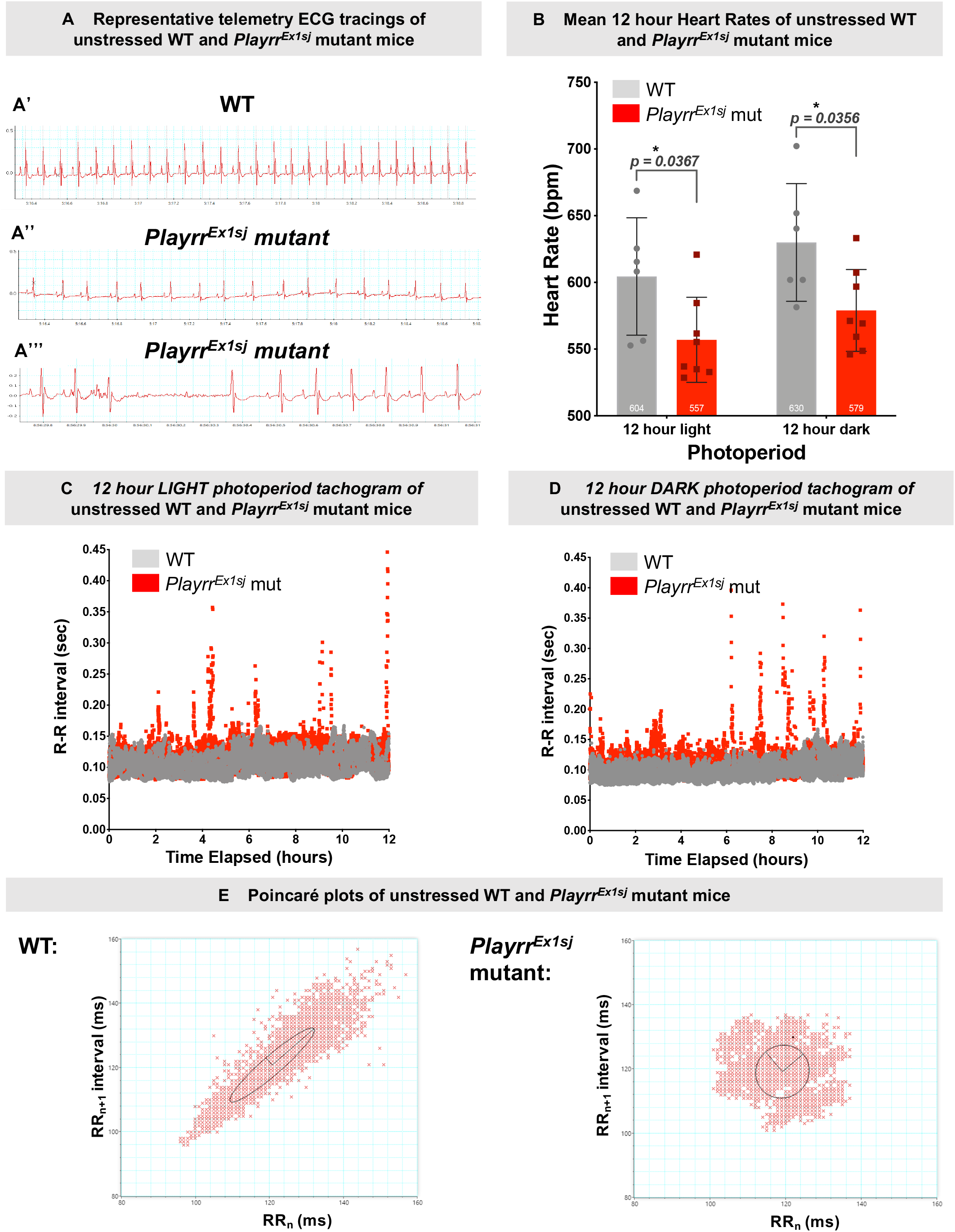
Heart rate and rhythm in 24-hour telemetry ECG recording in unstressed adult *Playrr*^*Ex1sj*^ mutant mice. **(A)** Representative telemetry ECG tracings of: (**A’**) normal sinus rhythm in WT (n= 6) and (**A’’**) irregular RR and bradycardia in *Playrr*^*Ex1sj*^ mutants (n=8). (**A’’’**) 1^st^ and 2^nd^ degree AV block observed in one *Playrr*^*Ex1sj*^ mutant. **(B)** Mean heart rate (bpm) by 12-hour photoperiod and 24-hour average in WT vs mutant mice. **(C)** 12-hour light (less active) photoperiod tachogram of unstressed WT (n=6) and *Playrr*^*Ex1sj*^ mutant (n=8) mutant mice. **(D)** 12-hour dark (active) photoperiod tachogram of unstressed WT (n=6) and *Playrr*^*Ex1sj*^ mutant (n=8) mutant mice. **(E)** Representative Poincare plots plotting subsequent (RRn+1) interval as a function of previous R-R interval (RRn) in 5000 consecutive beats *Playrr*^*Ex1sj*^ mutant mouse (right panel) and WT control (left panel).

### Loss of *Playrr* has no effect on SAN patterning

To investigate how loss of *Playrr* affects cardiac conduction, we first asked whether loss of *Playrr* leads to perturbed SAN patterning. To this end, we used ISH to investigate *Pitx2* expression in the left and right SAN domains at E8.5 (**Fig. S1B**) and at E10.5 (**Fig. S1C**). In *Playrr*^*Ex1sj*^ mutant embryos (E8.5 n=3; E10.5 n=5), *Pitx2* expression remained restricted to the left SAN domain, left sinus venosus, and left atrial myocardium. Importantly, no ectopic *Pitx2* expression was detected in any of the right sided corresponding domains, indicating that loss of *Playrr* does not lead to a double-left isomerism (**Fig. S1BC**).

To corroborate these findings, we investigated the expression of *Hcn4 (*Hyperpolarization-activated cyclic nucleotide-gated channel 4), a key marker of SAN (Stieber et al., 2003) in *Playrr*^*Ex1sj*^ mutant mice using the transgenic reporter allele, *Hcn4-CatCH*-IRES-*LacZ*.^30^ In both WT and *Playrr*^*Ex1sj*^ mutant *Hcn4-CatCH-*IRES-*LacZ* transgenic embryos, patterns of LacZ staining are similarly found in the primary heart tube (left ventricle, first heart field), sinus venosus, LR venous myocardium, and in the right sided SAN, with no corresponding expression on the left (n=3) **(Fig. S1D)**. These data suggest that loss of *Playrr* affects cardiac conduction independent of LR SAN patterning.

### Loss of *Playrr* is associated with an altered atrial substrate in adult mice

Previous work suggests that Connexin 43 (Cx43), a key gap junction protein that regulates cardiac rhythm, may be indirectly regulated by Pitx2, via interactions upstream of Shox2 and Tbx3.^9^ We therefore hypothesized that mis-localization of Cx43 within the atria of *Playrr*^*Ex1sj*^ adult mice could provide a substrate for the development of AF. Immunostaining revealed that Cx43 in WT left atria (LA) and right atria (RA) was primarily localized at the short ends of the myocytes, perpendicular to the myofibrils at the area of the intercalated disc (ID). Less frequently, Cx43 signal was also localized at the long edges of the myofibrils (lateralization), or in diffuse “pockets” within the WT cells (diffuse, DIF) (**Fig. 5A, D**). In contrast to the WT atria, Cx43 was primarily diffusely distributed within the cells in *Playrr*^*Ex1sj*^ mutant mice, with less IF signal noted at the area of the ID (**Fig. 5B, E**). These localization patterns were quantified in images from WT (LA, n=5; RA, n=5) and *Playrr*^*Ex1sj*^ mutant (LA, n=11; RA, n=11) samples. A significant decrease in Cx43 localization at the ID was noted in *Playrr*^*Ex1sj*^ mutant LA samples compared to WT LA controls (p=0.0459 for LA). While a similar significant change in Cx43 ID localization was not appreciated in the RA samples (p=0.467 for RA), we did note a significant increase in diffuse localization of Cx43 in *Playrr*^*Ex1sj*^ mutant RA samples (p=0.029), along with a trend towards increased diffuse Cx43 signal in *Playrr*^*Ex1sj*^ mutant LA samples. There was no significant difference in lateralization of Cx43 across the WT RA and LA samples (data not shown). Lastly, we quantified the ratio of diffuse (DIF)/ID signal in *Playrr*^*Ex1sj*^ mutant and WT LA and RA in order to further understand the contrasting localization patterns of Cx43 in the WT versus mutant samples. We found a statistically significant increase in *Playrr*^*Ex1sj*^ mutants for both LA (p=0.0002) and RA (p=0.0308) samples.

**Fig. 5.**
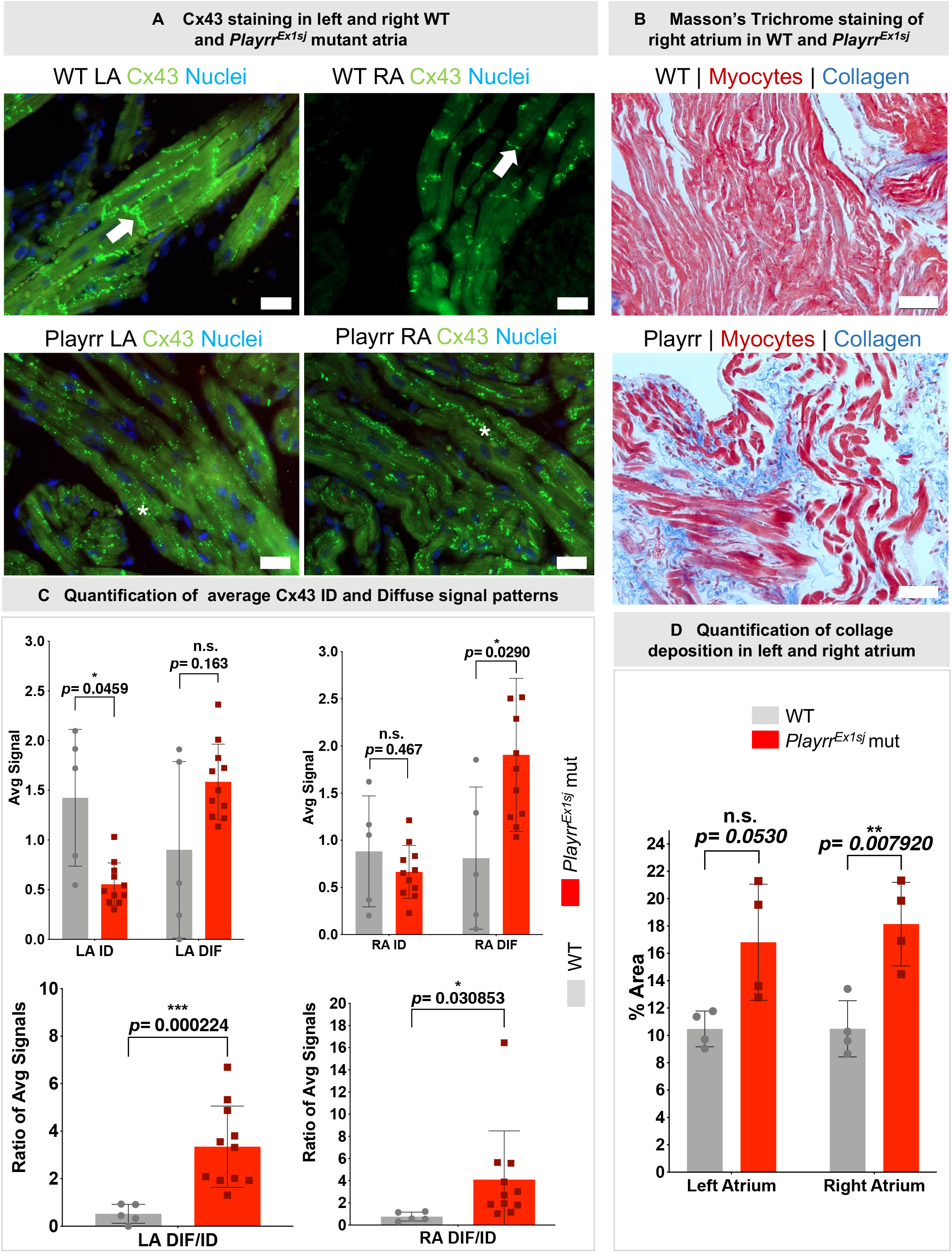
Localization of Connexin 43 and atrial fibrosis in adult Playrr^Ex1sj^ mutant mice. **(A)** Cx43 (green) immunofluorescent staining in WT left (left column) and right (right column) atrial myocytes from WT (top row) and *Playrr*^*Ex1sj*^ mutant (bottom row) adult mutant hearts. Signal at the short edges of the myocytes correspond to the intercalated disc (ID, white arrow). Cx43 diffuse pattern occurs within the cells (white asterisk). Scale bars = 20 μm. **(B)** Representative Masson’s trichrome stained sections of right atrium from WT (n=4, top) and in *Playrr*^*Ex1sj*^ mutants (n=4, bottom) adult mice. Myocytes are stained red and collagen deposition is stained blue. **(C)** Quantification of Cx43 localization patterns (ID or diffuse) in WT (grey) and *Playrr*^*Ex1sj*^ mutants (red) in left atrium (left column) and right atrium (right column). Ratio of diffuse/ID signal is depicted (bottom row). **(D)** Comparison of % area collagen deposition in LA and RA from WT (grey) and *Playrr*^*Ex1sj*^ mutant (red) sections.

We also considered whether an increase in fibrosis in RA and LA could provide a further substrate for the development of arrhythmias in *Playrr*^*Ex1sj*^ mutants. We found that collagen deposition, as quantified by analysis of Masson’s Trichrome staining, was significantly increased in *Playrr*^*Ex1sj*^ mutant RA compared to WT (p=0.00792). Although collagen deposition observed in *Playrr*^*Ex1sj*^ mutant LA was increased compared to WT, the result was not significant (p=0.0530). Collagen deposition comprised an average of 10.5% of the total area in WT sections of RA (n=4) and 18.1% in *Playrr*^*Ex1sj*^ mutant RA (n=4) (**Fig. 5C’’’**; p=0.007920). Taken together, mis-localization of Cx43 and increased fibrosis in *Playrr*^*Ex1sj*^ mutant mice may provide a substrate for the development of AF.

### Chromatin run-on and sequencing maps demonstrate changes in transcriptional landscape in *Playrr*^*Ex1sj*^ mutant hearts

To identify molecular pathways underlying arrhythmogenicity in adult *Playrr*^*Ex1sj*^ mutant mice, we performed Chromatin Run-On and Sequencing (ChRO-seq), which is able to measure the genome-wide abundance of RNA polymerase and nascent RNA transcription in solid tissue samples.^43^ By mapping the position and orientation of RNA polymerase I-III genome-wide, ChRO-seq provides insights into transcription of protein coding genes, noncoding RNAs, as well as the activity of regulatory elements such as enhancers.^43^ We used ChRO-seq to analyze pooled left and right atrial tissues from age-matched (32 weeks) *Playrr*^*Ex1sj*^ WT and *Playrr*^*Ex1sj*^ mutant mice (**Fig. 6A**; WT, n=4 vs *Playrr*^*Ex1sj*^ mutant, n=4). All samples were highly correlated, indicating that transcription was similar in the adult heart of all mice examined (**Fig. S2**). Principal component analysis (PCA) showed that WT and *Playrr*^*Ex1sj*^ mutant mice were separated on both PC1 and PC2, suggesting that transcriptional profiles of WT vs PlayrrEx1sj mutants cluster based on genotype and can be differentiated (**Fig. 6B**; WT, n=4 vs *Playrr*^*Ex1sj*^ mutant, n=4). Moreover, we identified 296 transcripts that had significant expression changes between WT and *Playrr*^*Ex1sj*^ mutant mice (**Fig. 6C**; FDR corrected p-value <0.01, DESeq2 ^43^; WT, n=4 vs *Playrr*^*Ex1sj*^ mutant, n=4). Gene ontology analysis of these 296 transcripts demonstrated strong enrichment in genes involved in heart development, ion transport, cell-cell adhesion, or motility (**Fig. 6E**). Notably, downstream targets of the transcription factor *Tbx5*, a key regulator of cardiac rhythm,^22^ were affected. Specifically, genes that have been previously linked to the development of atrial arrhythmias^52,53^ *Hcn1* and *Ryr2* (Ryanodine Receptor 2), the major Ca^2+^ channel protein in cardiomyocytes, showed 2- and 4-fold higher expression in *Playrr*^*Ex1sj*^ mutant atria, respectively (**Fig. 6C-D**). Altered expression of integral ion channels, possibly driven by overexpression of Tbx5, may play a key role in the development of AF and SND in *Playrr*^*Ex1sj*^ mutant mice. Indeed, misregulation of *Tbx5* has previously been linked to atrial arrhythmias by affecting the expression of *Pitx2* and several downstream ion channel genes.^22^ Therefore, we asked whether the *Tbx5* binding motif was enriched in enhancers showing evidence of higher Pol II loading in *Playrr*^*Ex1sj*^ mutant mice. ChRO-seq provides the location of enhancer-templated RNAs, which are accurate markers of both promoters and enhancers, collectively called transcription initiation regions (TIRs).^43^ We used dREG to identify the location of 52,257 candidate TIRs.^49^ One WT replicate (WT_R1) was excluded in this experiment, due to its transcriptional similarities to *Playrr*^*Ex1sj*^ mutants. After excluding this dataset, we identified 154 candidate TIRs that were more active in *Playrr*^*Ex1sj*^ mutant mice (FDR corrected p-value < 0.1, DESeq2). The motif that was most highly enriched in these 154 TIRs bore the AGGTGT core that is characteristic of all Tbx-family transcription factors, including *Tbx5* (**Fig. 6F**). Other motifs included forkhead-box transcription factors, NF-kB1 and Rfx family (**Fig. 6F**). Thus, our results suggest that concurrent with previous studies,^22,52^ misregulation of *Tbx5* may play a major role in downstream transcriptional changes in adult *Playrr*^*Ex1sj*^ mutant hearts with other Tbx-family transcription factors also contributing.

**Fig. 6:**
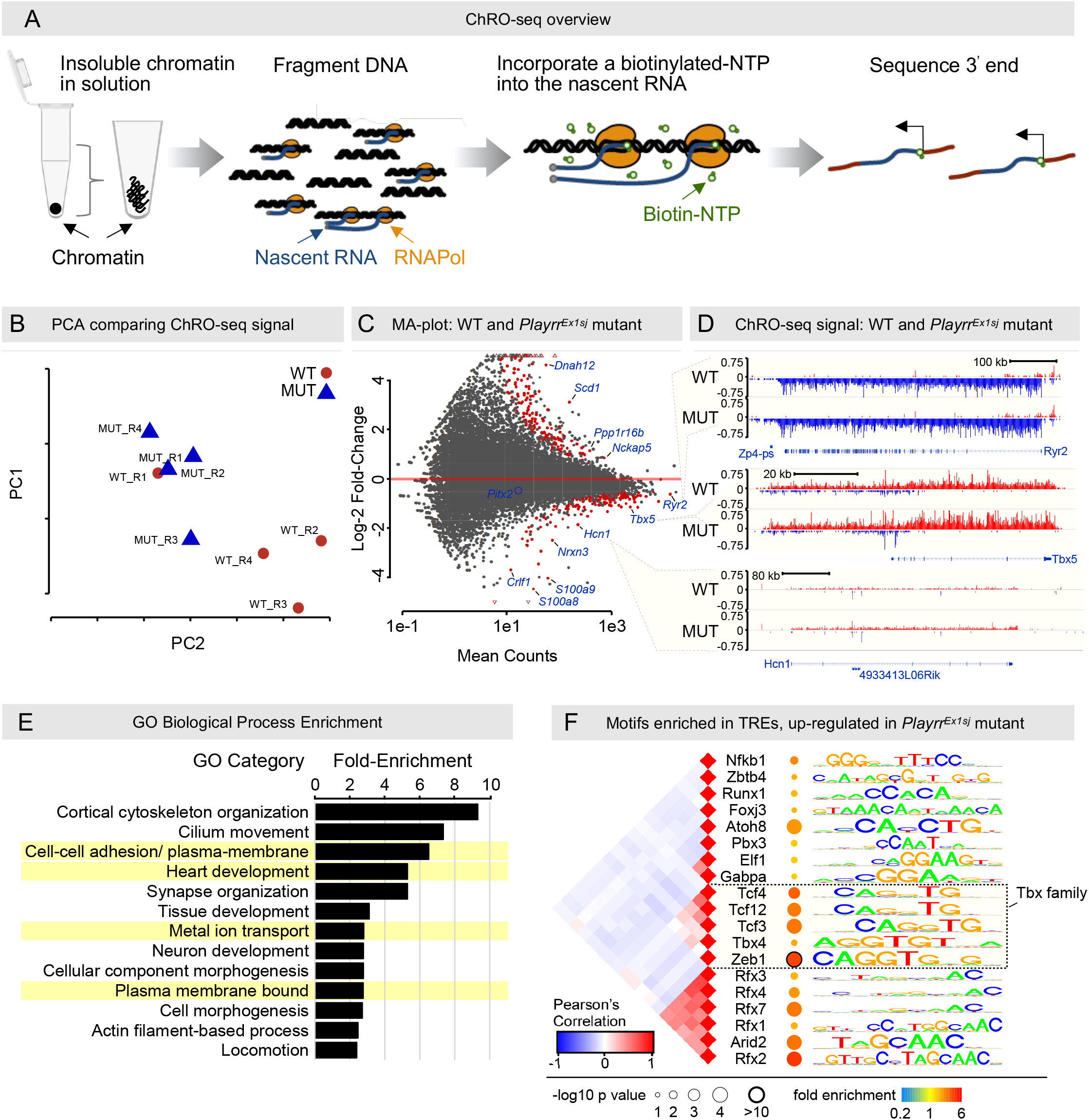
ChRO-seq transcriptomic analysis of *Playrr*^*Ex1sj*^ atrial tissue. **(A)** ChRO-seq measures primary transcription in isolated chromatin. Isolated chromatin from atrial tissue is resuspended into solution, incubated with biotinylated nucleotide triphosphate (NTP), purified by streptavidin beads, and sequenced from the 3’ end. The run-on was designed to incorporate a biotinylated NTP substrate into the existing nascent RNA that provides a high-affinity tag used to enrich nascent transcripts. The biotin group prevents the RNA polymerase from elongating after being incorporated into the 3’ end of the nascent RNA when performed in the absence of normal NTPs, thus enabling up to single-nucleotide resolution for the polymerase active site. **(B)** Principal Components Analysis (PCA) comparing ChRO-seq signal in WT (n=4; red circles) and *Playrr*^*Ex1sj*^ mutants (n=4; blue triangles). **(C)** MA plot visualizing gene expression in WT and *Playrr*^*Ex1sj*^ mutants. **(D)** ChRO-seq signal in WT and *Playrr*^*Ex1sj*^ mutants of *Ryr2* (top), *Tbx5* (middle), *and Hcn1* (bottom). **(E)** Bar chart of Gene Ontology (GO) Biological Process fold-enrichment in *Playrr*^*Ex1sj*^ mutants vs WT controls. **(F)** Transcription factor binding motifs enriched in TREs upregulated in *Playrr*^*Ex1sj*^. Pearson’s rank correlation heatmap (left) shows correlation in DNA binding sites matching each motif. Radius of the circle after the gene name represents the p value across samples, with color representing magnitude of enrichment (red) or depletion (blue). Tbx, T-box family.

**Fig. 7.**
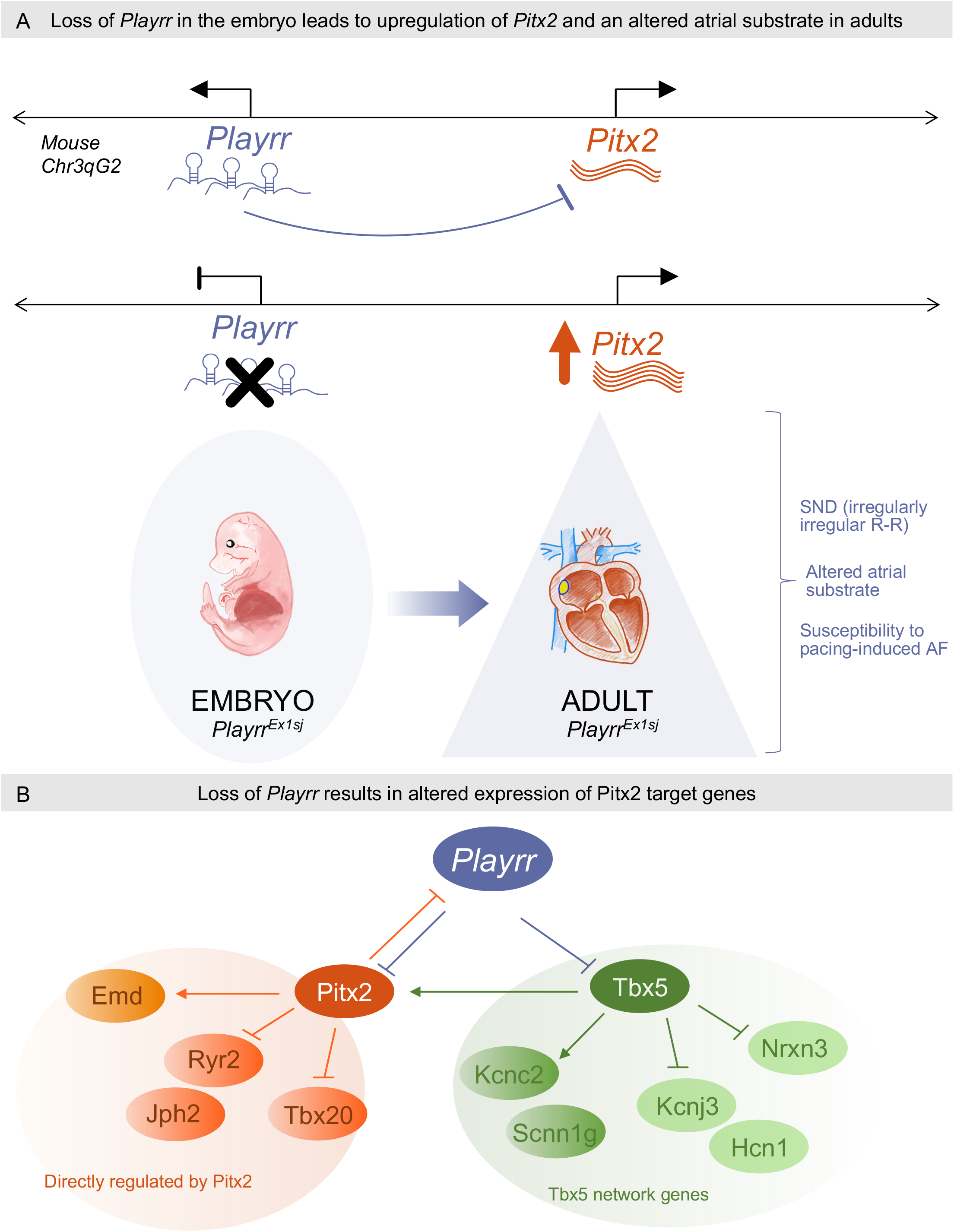
Model for the role of the lncRNA *Playrr* in cardiac arrhythmogenesis. **(A)** *Playrr*^*Ex1sj*^ mutant embryos at E14.5 have upregulation of *Pitx2c*, the predominant *Pitx2* cardiac isoform. Adult *Playrr*^*Ex1sj*^ mutants have an altered atrial substrate, evidence of sinus node dysfunction, and susceptibility to pacing-induced AF. **(B)** *Playrr*^*Ex1sj*^ mutant atria have altered expression of *Pitx2* target genes, including dysregulation of *Pitx2* and *Tbx5* network genes previously implicated in control of atrial substrate and/or atrial rhythm.

Collectively, we propose that loss of *Playrr* in the embryo leads to upregulation of *Pitx2* and *Tbx5* expression, dysregulated expression of core regulators of cardiac rhythm genes, and ultimately susceptibility to atrial arrhythmias in adult life.

## Discussion

Here we have utilized the CRISPR/Cas9-generated mutant mouse allele, *Playrr*^*Ex1sj*^ (Welsh et al., 2015), a splice-site mutation that selectively degrades the *Playrr* lncRNA transcript while leaving binding sites in the associated DNA enhancer (e926) intact, to investigate the genetic mechanisms of *Pitx2* linked arrhythmias. Specifically, we sought to identify the transcriptional and developmental functions of *Playrr*, an evolutionarily conserved lncRNA arising from noncoding DNA at the *Pitx2* genomic locus. In doing so, we aimed to investigate the mechanisms by which noncoding regulatory elements at the *Pitx2* locus may mediate transcriptional regulation of *Pitx2* necessary for cardiac function.

### AliveCor as a novel screening tool for arrhythmias

Using both AliveCor^®^ surface ECG screening and telemetry ECG, we demonstrate that *Playrr*^*Ex1sj*^ mutant mice have resting cardiac arrhythmias evident of SND, a major predisposing risk factor for AF. To our knowledge, this is the first study utilizing the AliveCor^®^ device for use in laboratory mice. The independent experimental validation of this device for arrhythmia screening in mice was concurrently performed in collaboration with colleagues in the Cornell University Center for Animal Resources and Education.^50^ Both AliveCor^®^ ECG screening on awake, restrained mice and 24-hour telemetry ECG in non-stressed mice under physiological conditions demonstrate that *Playrr*^*Ex1sj*^ mutants have bradycardia and increased beat-to-beat interval variability, compared to controls. Importantly, these signs represent diagnostic criteria for SND, an important arrhythmia and risk factor for AF.

### *Playrr* vs *Pitx2* specific spatiotemporal gene expression during cardiac development

We discovered that *Playrr* is expressed in a right-specific pattern in cardiac tissue opposite the left-specific *Pitx2*, in the primordial SAN domain. We demonstrate specific spatiotemporal expression of *Playrr* in the heart, notably in the previously documented embryonic domain of the SAN,^14,54,55^ a small restricted mesenchymal region in the right pleuropericardial fold at the junction of the right atrium and the right common cardinal vein at E10.5. This is in contrast to *Pitx2*, which in addition to being expressed in the corresponding left SAN domain, demonstrates much broader expression dorsally and ventrally in all left splanchnopleure derived tissues (**Fig. 2B**). Computational modeling approaches have been developed to infer biological functions of lncRNAs from their highly tissue restricted expression.^56,57^ Our findings in this study reinforce that the expression pattern of a lncRNA, like that of a protein coding gene, may be used to predict its role in biological and pathophysiological processes at the organismal level.^57^

### Loss of *Playrr* in the embryo is linked with later onset of SND in mature adult mice

Our telemetry ECG studies in 28-week-old adult *Playrr*^*Ex1sj*^ mutant mice provide the first evidence that embryonic loss of *Playrr* leads to SND. Similar cardiac arrhythmias were previously characterized in mice with germline deficiency in *Pitx2-*microRNA pathways (cardiac specific *miR-17-92* knockout),^51^ conditional knockout of *Pitx2* in postnatal atrium,^8^ and *Hcn1* knockout mice.^26^ Thus, it is tempting to speculate that *Playrr* may be involved in Pitx2-mediated cardiac function. Unlike in 28-week-old *Playrr*^*Ex1sj*^ mutant mice, telemetric recordings in 12-week-old *Playrr*^*Ex1sj*^ mutants did not show significant differences in HR or RR intervals compared to WT controls (data not shown), suggesting that embryonic loss of *Playrr* results in progressive tissue changes that lead to the development of SND later in mature, middle-aged adults. Our findings suggest that unlike previous genetic mouse models that develop juvenile and early adult-onset arrhythmias,^26,58^ *Playrr*^*Ex1sj*^ mutant mice may develop SND and predisposition to AF with aging, mirroring SND onset and increased risk of paroxysmal AF in otherwise healthy aging adult humans. Indeed, *Playrr*^*Ex1sj*^ mutant mice are predisposed to pacing-induced AF, at a similar frequency to that found in *Pitx2* heterozygous mice,^9^ suggesting that loss of *Playrr* in the embryo has lasting transcriptional consequences in the adult heart. Moreover, while increased *Pitx2c* was detected in the *Playrr*^*Ex1sj*^ mutant embryos at E14.5, its upregulation was not detected in adult heart samples via our ChRO-seq data set. *Pitx2* expression in the adult heart is very low,^9^ and small changes – if present – may be below the detection level for ChRO-seq. Minute, cell – specific increases in *Pitx2* transcription in adult *Playrr*^*Ex1sj*^ mutant hearts (below the detection level for ChRO-seq) may lead to large differences in downstream expression of transcription factors and their cell effectors. Alternatively, embryonic increase in *Pitx2* expression may result in lasting transcriptional changes which persist in the adult heart.

### Atrial substrate is perturbed with mis-localization of Cx43 and increased fibrosis

Adult *Playrr*^*Ex1sj*^ mutant mice show evidence of an altered atrial substrate, consisting of mis-localization of Cx43 and increased collagen deposition in RA and LA. These abnormalities suggest an altered myocardial landscape that can potentiate both SND and AF.^59,60^ While the molecular mechanisms by which loss of *Playrr* results in altered atrial substrate remain unclear, our data suggest that loss of *Playrr* negatively impacts *Pitx2* regulation of key transcriptional events in the heart during development. Indeed, *Pitx2* is a known upstream transcriptional regulator of *Cx43*, via *Shox2* regulation of *Tbx3*.^*54,61*^ Furthermore, our ChRO-seq data reveal a significant increase in *Tbx5* transcription, a direct transcriptional regulator of *Ryr2, Hcn1*, and *Cx40* (*Gja5*), an integral atrial gap junction protein. The relationship between Cx40 and Cx43 in atrial myocytes has been previously investigated,^62^ and disruption of the complex interactions between different connexin isomers within the atrial ID may provide a substrate for atrial arrhythmias.^63,64^

Despite significant mis-localization of Cx43 in the atria of *Playrr*^*Ex1sj*^ mutant mice, ChRO-seq analysis did not identify a significant differential *Cx43* mRNA expression compared to age-matched WT mice. This is consistent with previous studies suggesting that Cx43 mis-localization may occur secondary to the cellular microenvironment independent of gene expression.^59,60^ We hypothesize that Cx43 mis-localization may be an indirect effect of other *Playrr*- and/or *Pitx2*-regulated changes to the cellular architecture, rather than altered transcriptional regulation.

Atrial sections from *Playrr*^*Ex1sj*^ mutant mice revealed increased collagen deposition when compared to WT counterparts. This is supported by changes in gene expression noted in our ChRO-seq data, where genes associated with cortical cytoskeleton organization and cell-cell adhesion, such as *Emd*, are significantly enriched in *Playrr*^*Ex1sj*^ mutant hearts compared to WT controls. Whether increased transcription of these genes is a direct result of loss of *Playrr*, or secondary to a cascade of events, including remodeling of ID proteins such as Cx43, remains to be determined.

Alterations of atrial substrate were noted in *Playrr*^*Ex1sj*^ mutant samples from both LA and RA, however, the findings varied between LA vs RA samples. Increased mis-localization of Cx43 at the ID in *Playrr*^*Ex1sj*^ mutant vs WT atria was observed in LA at the ID and for diffuse signal in the RA. A significant increase in collagen deposition was observed in RA samples but was not significant in LA collagen deposition was observed in *Playrr*^*Ex1sj*^ mutants compared to age-matched controls. Further research is necessary to determine whether loss of *Playrr* preferentially affects specific right- or left-sided pathways involved in the development of fibrosis, and/or mis-localization of Cx43 in the atria.

### ChRO-seq reveals overexpression of *Tbx5*-directed regulatory network and dysregulation of ion channels

Altered expression of genes integral to maintaining the normal cardiac rhythm was evaluated using ChRO-seq. These analyses revealed additional transcriptional changes in genes linked to the development of cardiac arrhythmias in *Playrr*^*Ex1sj*^ mutant mice. In addition to upregulation of *Tbx5*, upregulation of *Hcn1* was noted. These genes were previously implicated in the generation of normal cardiac rhythm.^22,58^ Notably, both *Pitx2*-dependent and *Pitx2-* independent transcriptional regulatory networks have been linked to control of LR atrial patterning, cardiac conduction, and atrial arrhythmias.^65^ *Tbx3*, via *Shox2* upregulation, promotes SAN phenotype and inhibits working atrial phenotype by inhibiting *Cx43*.^15,54,61^ Misregulation of the Tbx family of genes in *Playrr*^*Ex1sj*^ mutants may result in alterations in electrical make-up of SAN (resulting in SND) and affect electrical coupling of working atrial myocytes, further predisposing to AF.

Of further interest, significant alterations in calcium handling genes were appreciated in *Playrr*^*Ex1sj*^ mutant mice. Both *Ryr2* and cardiac *Casq2* (Calsequestrin 2), the major sarcoplasmic reticulum calcium-binding protein, were significantly upregulated in *Playrr*^*Ex1sj*^ mutants (p<0.001 and p<0.02, respectively) compared to WT controls.^66^ We also observed an upregulation of *ATP2b2*, the plasma membrane calcium transporting ATPase 2, and the genes encoding several calcium channels, including KCNJ3 (Potassium Inwardly Rectifying Channel Subfamily J Member 3). Of particular importance to the SAN is the spontaneous depolarization (automaticity) that sets the normal cardiac rhythm, and which is partially regulated by the calcium clock.^67^ The calcium clock refers to the spontaneous release of calcium from the sarcoplasmic reticulum and more specifically from *Ryr2* that span the sarcoplasmic reticulum membrane.^67^ Importantly, inhibition of the Ryr2 protein results in slowed pacemaker activity^68^ and human patients with mutations in *RYR2* or *CASQ2* display episodes of catecholamine-induced polymorphic ventricular tachycardia in response to physical or emotional stress^69^ and are at increased risk for SND, AF, AV node dysfunction, and atrial standstill.^16,70^ Finally, a gain-of-function mutations in *KCNJ3* results in SND and AF.^71^ Thus, misregulation of integral calcium handling genes may contribute to the development of SND and AF in *Playrr*^*Ex1sj*^ mutant mice. *Pitx2* dosage is fundamentally linked to asymmetric organ development and different organs have distinct requirements for *Pitx2* dosage.^72,73^ We found significant ∼1.5-fold upregulation of the *Pitx2c* isoform and increased (non-significant, p=0.09) *Pitx2ab* isoforms in *Playrr*^*Ex1sj*^ mutant mice at E14.5 in pooled organ primordia (heart, lungs, intestines). These findings suggest that *Pitx2c* expression may be altered in *Playrr*^*Ex1sj*^ mutant mice during embryonic heart development and this may predispose to development of AF in adult life. Importantly, although the work of several key groups has implicated levels of *Pitx2* dosage in regulating SAN patterning, our analyses using ISH show conserved LR *Pitx2* expression and patterning of the embryonic SAN in *Playrr*^*Ex1sj*^ mutant embryos (E8.5, E10.5) (**Fig. S1)**. These data suggest that loss of *Playrr* predisposes to AF independent of SAN patterning, implicating a role for *Playrr* in cardiac morphogenesis and electrophysiology. Consistent with this hypothesis, *Pitx2* dosage regulates not only SAN developmental genes (*Shox2, Hcn4, Tbx3*),^9^ but also the expression of genes that direct cardiac electrophysiology.^8^ For example, atrial-specific conditional Pitx2 mouse mutants, in which right-sided SAN formation is intact, supports a role of *Pitx2* impairment in cardiac electrophysiology.^10^ Interestingly, *KCNN3*, which has recently been linked to AF by GWAS,^6,17^ is also altered by *Pitx2* insufficiency.^74^ Further evidence of the dysregulation of cardiac electrophysiology by *Pitx2* impairment has also been reported using mRNA microarray analyses in *Pitx2* heterozygous adult mutant mice.^7^ Collectively, our findings suggest a role for *Playrr* in the heart conduction system and support the hypothesis that dysregulation of *Playrr* expression can be responsible for susceptibility to AF.

In summary, our results reveal a new role for the *Playrr* lncRNA in regulating Pitx2c dosage and susceptibility to AF. We show a role for *Playrr* in regulating *Pitx2c* dosage, which may lead to misregulation of genes downstream of *Pitx2* and/or *Tbx5* and resulting atrial arrhythmias. Having the tools and in vivo mouse models to investigate the functional relevance of these noncoding elements represents a critical opportunity to identifying mechanistic links between transcriptional regulation of *Pitx2*, genomic mechanisms modulated by lncRNAs, and disease pathogenesis of cardiac arrhythmias. Future studies investigating the status of these critical downstream transcriptional targets of Pitx2 in *Playrr*^*Ex1sj*^ mutant mice, both during development and in the adult, will lend invaluable insight into the role of this functional lncRNA within gene regulatory networks relevant to cardiac disease. Limitations notwithstanding, our findings suggest a role for the lncRNA, *Playrr*, in regulating the molecular make-up and electrical functions of both the sinus node and working atria, and may help to explain the “missing link” of how sinus node dysfunction predisposes some cohorts to the development of AF.

## Abbreviations

AF: atrial fibrillation
GWAS: genome-wide association study
lncRNA: long noncoding RNA
ISH: in situ hybridization
qRT-PCR: quantitative real time polymerase chain reaction
SAN: sinoatrial node
SND: sinus node dysfunction
SNP: single nucleotide polymorphism
ChRO-seq: chromatin run-on & sequencing

## Acknowledgements

We thank Dr. Frank Lee and Dr. Michael Kotlikoff of CHROMus/Kotlikoff lab for providing *Hcn4-CatCH-IRES-LacZ* transgenic mice; Dr. Marcos Simoes-Costa for guidance and reference for adapting TaqMan Cells-to-Ct kit for qRT-PCR on low input, microdissected organs from single embryos; Rachael Labbitt and Scott Butler for help with telemetry implant surgeries; Joseph Choi for technical assistance. Dr. Manuel Martin-Flores for providing access to LabChart Pro software. We are grateful to all members of the Kurpios lab for feedback on the manuscript.

## Sources of Funding

NIH F30OD021454

NIH NIDDK R01 DK092776

NIH NIDDK R01 DK107634

NIH NHGRI R01 HG009309

March of Dimes – 1-FY11-520

## Author Contributions

Conceptualization: F.LC., E.M.O., and N.A.K. Methodology: F.L.C., E.M.O, S.C., N.L. and N.A.K.

Investigation: F.L.C., E.M.O., S.C, C.G.D, N.L., J.P.L, S.K.P., B.S., C.C., S.A.K, R.D., E.K.D

Writing – original draft: F.L.C.,

Writing – review & editing: F.L.C., E.M.O, J.FM. and N.A.K. Funding acquisition: F.L.C. and N.A.K.;

Supervision: E.M.O., C.G.D, J.F.M., and N.A.K.

## SUPPLEMENTAL FIGURE LEGENDS

**Fig. S1.**
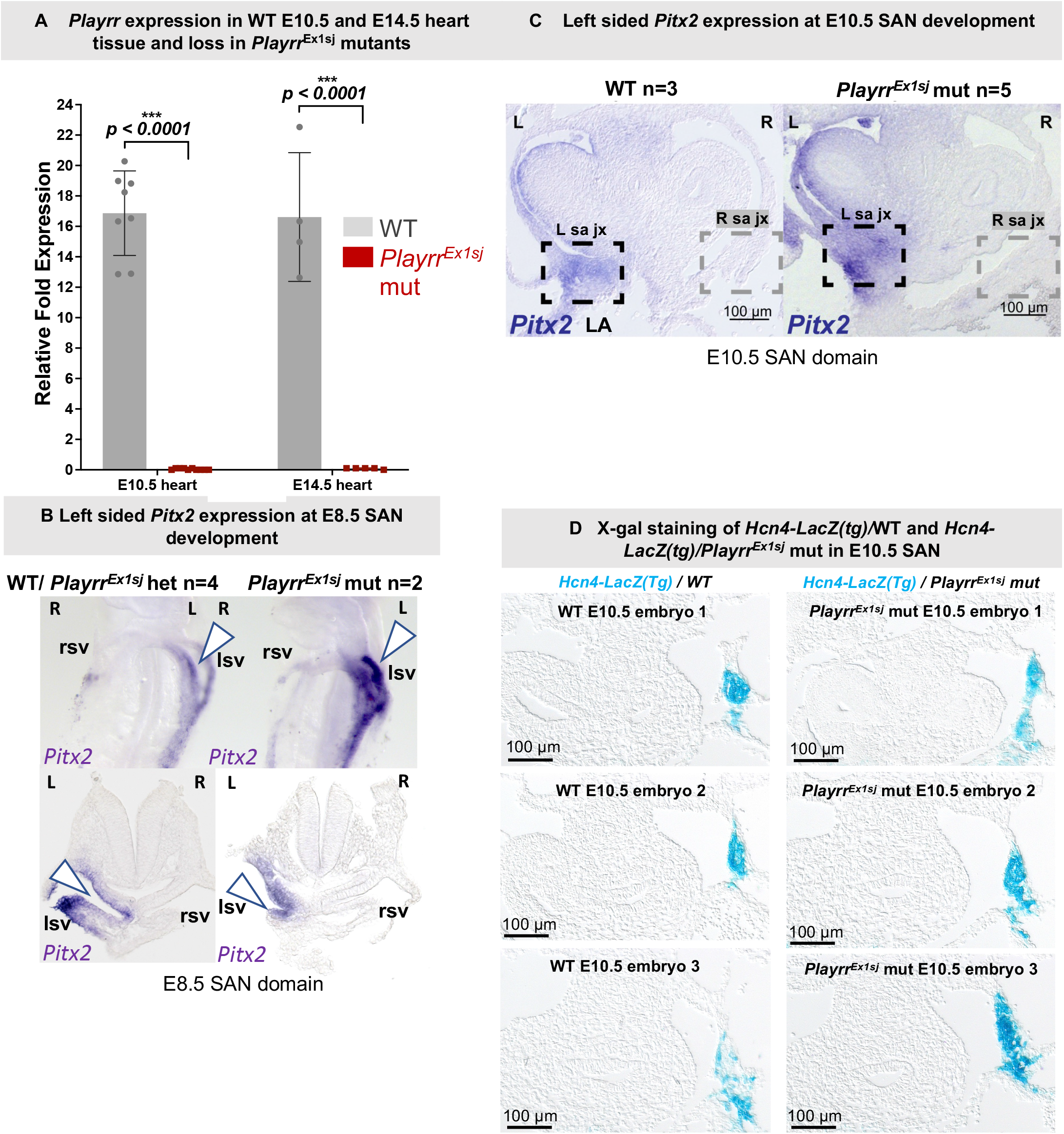
LR pattern of *Pitx2* expression is unchanged in SAN domain of Playrr^Ex1sj^ mice. **(A)** *Playrr* transcript assayed by qRT-PCR in microdissected E10.5 whole hearts (left) and E14.5 whole hearts (right). WT depicted in grey bars; *Playrr*^*Ex1sj*^ mutants in red bars. **(B)** Left sided *Pitx2* expression in WT or *Playrr*^*Ex1sj*^ het controls (n=4) (left) vs mutant (n=3) embryos at E8.5. Ventral view of whole mount embryos is shown in the top panels; sections depicted in the bottom panels. **(C)** *Pitx2* expression in sectioned whole mount ISH assayed in WT (n=3) and *Playrr*^*Ex1sj*^ mutant (n=5) embryos. Black dashed box demarcates left sided SAN domain; grey dashed box demarcates right sided SAN domain. **(D)** X-gal staining of *Hcn4-LacZ transgenic (Tg)/ Playrr*^*Ex1sj*^ WT (n=3) vs *Hcn4-LacZ transgenic (Tg)/ Playrr*^*Ex1sj*^ mutant (n=3) E10.5 embryos. Black dashed box highlights right sided SAN cells expressing *Hcn4-LacZ*. Representative sections from three different embryos of each genotype are shown.

**Fig. S2.**
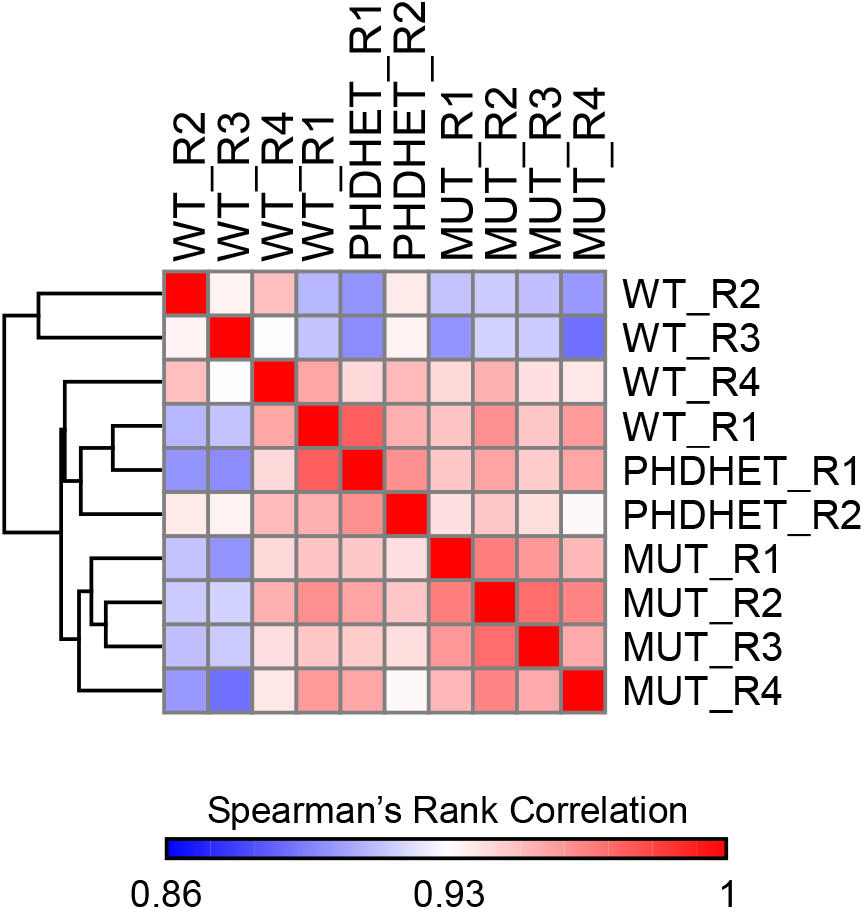
Spearman’s Rank Correlation matrix. Spearman’s Rank Correlation matrix from ChRO-seq analysis of WT (n=4) vs *Playrr*^*Ex1sj*^ mutant (MUT; n=4) adult hearts.

